# Single-cell transcriptomic and proteomic analysis of Parkinson’s disease Brains

**DOI:** 10.1101/2022.02.14.480397

**Authors:** Biqing Zhu, Jae-Min Park, Sarah Coffey, I-Uen Hsu, TuKiet T. Lam, Pallavi P. Gopal, Stephen D. Ginsberg, Jiawei Wang, Chang Su, Hongyu Zhao, David A. Hafler, Sreeganga S. Chandra, Le Zhang

## Abstract

Parkinson’s disease (PD) is a prevalent neurodegenerative disorder where recent evidence suggests pathogenesis may be mediated by inflammatory processes. The molecular architecture of the disease remains to be fully elucidated. We performed single-nucleus transcriptomics and unbiased proteomics using postmortem tissue obtained from the prefrontal cortex of 12 individuals with late-stage PD and age-matched controls. We analyzed ∼80,000 nuclei and identified eight major cell types, including brain-resident T cells, each with distinct transcriptional changes in line with the known genetics of PD. By analyzing Lewy body pathology in the same postmortem tissue, we found that α-synuclein pathology is inversely correlated with chaperone expression in excitatory neurons. Examining cell-cell interactions, we found a selective abatement of neuron-astrocyte interactions and enhanced neuroinflammation. Proteomic analyses of the same brains identified synaptic proteins in prefrontal cortex that were preferentially downregulated in PD. Strikingly, comparing this dataset to a regionally similar published analysis for Alzheimer’s disease (AD), we found no common differentially expressed genes in neurons, but identified many shared differentially expressed genes in glial cells, suggesting that disease etiology in PD and AD are likely distinct. These data are presented as a resource for interrogating the molecular and cellular basis of PD and other neurodegenerative diseases.

**One Sentence Summary:** We provide an extensive single cell analysis profiling nearly 80,000 brain nuclei from prefrontal cortex of late-stage Parkinson’s disease brains, demonstrate that α-synuclein pathology is inversely correlated with chaperone expression in excitatory neurons, found a selective abatement of neuron-astrocyte interactions with enhanced neuroinflammation, and augmented the study with proteomic analysis and cross-comparisons with Alzheimer’s disease datasets, providing valuable insights into the pathways of neurodegeneration and a deep definition of the underlying molecular pathology for Parkinson’s disease.

## INTRODUCTION

Parkinson’s disease (PD) is a common neurodegenerative movement disorder that impacts approximately 1 million people in the USA and nearly 6 million worldwide(*1, 2*). The neuropathological hallmarks of PD are loss of substantia nigra neurons and intracellular α-synuclein aggregates known as Lewy bodies. Sporadic/idiopathic PD comprises 90% of cases, indicating that the PD phenotype can arsie from varied, complex interactions of environmental and genetic risk factors. Several mechanisms have been implicated in the pathogenesis of familial PD syndromes, including α-synuclein proteostasis deficits, mitochondrial dysfunction, and synaptic vesicle cycling defects. More recently, neuroinflammatory pathways involving glial and immune cells have been suggested to contribute to PD pathogenesis(*3*). However, whether these pathways contribute to sporadic disease remains to be clarified. Moreover, the relative contributions of neuronal versus glial cell populations to PD pathophysiology remains unknown. Single cell and single nucleus RNA sequencing technologies have recently become robust and rigorous methods for assessing the heterogeneity of cell types in various diseases(*4*). While single cell technologies including multiomics analysis are becoming popular methods to the study of neurodegeneration, these tools are only now beginning to be applied to PD research, especially employing postmortem human brain tissues(*5*).

To obtain an accurate and unbiased assessment of the complex cellular changes associated with PD, we performed single cell transcriptomics and proteomics using late-stage PD brains and age-matched controls. We investigated the prefrontal cortex, as this region exhibits Lewy body pathology in late-stage PD(*6*), and because bulk transcriptomic analysis of this region in PD brains indicated the need for cell-type resolution(*7*). By focusing on the prefrontal cortex, we can also make robust cross-comparisons between PD and Alzheimer’s disease (AD) based on published AD datasets for this brain region(*8*). To our knowledge, the datasets presented herein represent the first single-cell portrait of PD brain prefrontal cortex. Taken together, our data reveal a unique cellular-level view of transcriptional and cortical protein changes associated with PD. Furthermore, we provide strong evidence for the presence of discrete Lewy body associated signatures, as well as revealing features in the PD dataset that share commonalities with other neurodegenerative diseases.

## RESULTS

### Single nucleus transcriptomic profiling of postmortem PD brains

To understand cellular diversity and disease-related transcriptional changes in PD, we performed single nucleus RNA sequencing (snRNA-seq) to profile postmortem human brain tissue from the prefrontal cortex Brodmann area 9 (BA9, Dorsolateral prefrontal cortex) of six PD patients and six sex- and age-matched healthy controls (mean age 75.9, range 63-96 years, Supplementary Table 1). We optimized an unbiased isolation protocol for human brain nuclei using sucrose gradient ultracentrifugation (Supplementary Fig. 1A-C, see Methods) followed by snRNA-seq on the 10x Chromium platform (10x Genomics). We profiled 77,384 brain nuclei and detected a median of 2,598 genes and 5,639 transcripts per nucleus (Supplementary Fig. 1D-F), after quality control filtering. To classify the major cell types in prefrontal cortex, we clustered all nuclei jointly across the 12 individuals (both PD and healthy controls) to produce 12 transcriptionally distinct clusters with highly consistent expression patterns across individuals (Fig. 1A-C). We identified and annotated eight major cell types with known marker genes, namely: excitatory neurons (ExN, *SLC17A7*) with three subclusters (ExN1, ExN2, ExN3), inhibitory neurons (InN, *GAD1*) with three subclusters (InN1, InN2, InN3), astrocytes (Astro: *AQP4*), microglia (MG: *ITGAM*), oligodendrocytes (Oligo: *MBP*), oligodendrocyte precursor cells (OPC: *PDGFRA*), endothelial cells (Endo: *CLDN5*), and brain-resident T cells (T cells: *CD3E*) (Fig. 1A, Supplementary Fig. 1H). We found unique or enriched RNA markers for each of the cell types, including *CUX2, CBLN2, HS6ST3* for excitatory neuron subcluster 1; *TSHZ2, POU6F2, FOXP2* for excitatory neuron subcluster 2; *NRG1, NEFL, CADPS2* for excitatory neuron subcluster 3; *ADARB2, GALNTL6, RGS12* for inhibitory neuron subcluster 1; *NXPH1, KIAA1217, SOX6* for inhibitory neuron subcluster 2; *FGF13, TMEM132D, UNC13C* for inhibitory neuron subcluster 3; *ST18, PLP1, MBP* for oligodendrocytes; *AQP4, GFAP, RYR3* for astrocytes; *SPP1, RUNX1, DOCK8* for microglia; *LHFPL3, TNR, MEGF11* for oligodendrocyte precursor cells; *EBF1, ADGRF5, TFPI* for endothelial cells; *SKAP1, GNLY, CD96* for brain-resident T cells (Fig. 1D). The relative proportions of five of the major cell types were roughly similar between PD and healthy controls, with the exceptions of oligodendrocytes, microglia, and T cells. We identified more microglia (healthy controls, n = 1699 nuclei; PD, n = 3684 nuclei; P = 0.09) and significantly increased brain resident T cells (healthy controls, n = 33 nuclei; PD, n = 243 nuclei; P = 0.02) in PD samples, indicating the presence of neuroinflammation (Fig. 1E, Supplementary Fig. 1G).

**Fig. 1.**
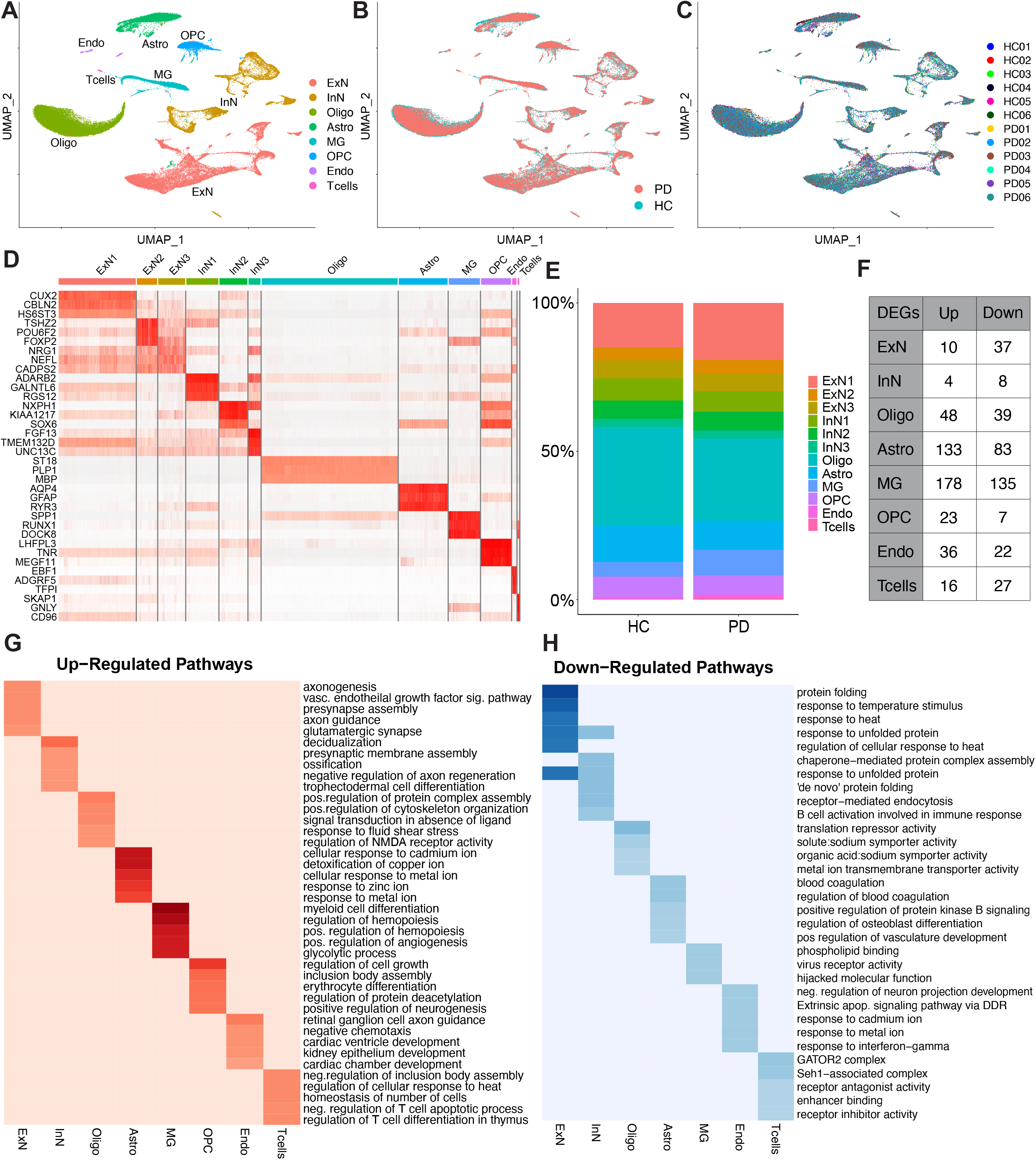
Single nucleus transcriptomic analysis of human brain prefrontal cortex reveals cell-type-specific changes in PD. (**A-C**), UMAP plotting of human brain nuclei (n = 77,384 nuclei), colored by (A) major human brain cell types: excitatory (ExN) and inhibitory (InN) neurons, oligodendrocytes (Oligo), astrocytes (Astro), microglia (MG), oligodendrocyte precursor cells (OPC), endothelial cells (Endo) and brain-resident T cells (Tcells); (B) disease diagnosis of either Parkinson’s disease (PD) or healthy controls (HC); or (C) individuals (from 12 individuals of 6 PD and 6 HC). (**D**) Heatmap of top marker genes in each brain cell type. (**E**) Brain cell type proportion grouped by disease diagnosis of PD or HC. (**F**) Differentially expressed genes (DEGs) counts for each brain cell type (log fold change > 0.75 for T cells; log fold change > 0.25, FDR corrected P-value adjusted < 0.05 for all the other cell types). (**G-H**) Gene ontology (GO) pathway analysis of DEGs between PD and HC for each brain cell type for either up-regulated pathways (G) or down-regulated pathways (H).

We next examined the cell type-specific gene expression changes between PD and healthy control prefrontal cortex and identified 806 unique differentially expressed genes (DEGs). Assigning the DEGs to the major cell types, we found that neurons showed fewer significant changes than glial cells, with astrocytes and microglia having the greatest number of DEGs (Fig. 1F, Supplementary Table 2). Both excitatory and inhibitory neurons revealed a signature of gene repression with 79% and 67% of downregulated DEGs, respectively. Conversely, all glial cells populations revealed an overall upregulation of gene expression with astrocyte and microglial changes being the most prominent.

We performed Gene Ontology (GO) pathway analysis of both downregulated and upregulated DEGs in each cell type (Fig. 1G-H, Supplementary Table 2). The most significantly upregulated pathway in neurons (Fig. 1G) consisted of GO terms associated with axonogenesis (*EPHA6, ROBO1, GAB1, PARD3*) and presynaptic assembly (*NLGN4Y, CBLN2*), suggesting compensatory synaptogenesis and synaptic alterations. We validated our snRNA-seq dataset using RNAscope *in situ* hybridization on an additional cohort of male PD and control brain prefrontal cortex sections for a few differentially expressed genes, such as *EPHA6* and *CBLN2* (Supplementary Fig. 3A) and quantified the RNAscope signals between PD and controls using QuPath (Supplementary Fig. 3A-B).

Upregulated pathways in oligodendrocytes indicate protein complex assembly (*HSPA1A, HSPA1B, FMN1*) and cytoskeleton organization (*P2RX7, RAPGEF3, FMN1, FCHSD2*). Interestingly, astrocytes showed enhanced expression of pathways related to detoxification of heavy metals (*MT1G, MT1F, MT3*), consistent with iron and other heavy metal deposition in PD brains(*9, 10*). Microglia in PD brains were enriched for the pathways of myeloid cell differentiation (*FOS, STAT3, RUNX1*), regulation of hemopoiesis (*BCL6, IL4R, JAK3*), and glycolytic process (*NFKBIZ, IL15, NFKBID*), all markers of microglia activation in response to disease. In OPC, regulation of cell growth (*VEGFA, HSPA1, HSPA1B*) and positive neurogenesis (*ZNF365, SYT1, MAPT*) pathways were observed displaying presumed regenerative responses toward neuroinflammation. As we unbiasedly profiled all brain nuclei without enrichment of immune cell nuclei, there were relatively few T cells present. Nevertheless, there was an enrichment of genes in brain-resident T cells in PD brains for pathways regulating homeostasis of cell number (*IL7R, IL15, MALT1*) and regulation of T cell differentiation (*CAMK4, CD28, ZAP70*), suggesting a prominent T cell response is present in PD, consistent with the newly discovered roles for T cells in PD(*11, 12*).

The most prominent downregulated pathways were in neurons (Fig. 1H). Protein folding and unfolded protein response pathways (*HSPB1, HSP90AB1, DNAJB6, HSPA1A, HSPH1*) were the most downregulated pathways within excitatory neurons in PD, signifying decreased chaperone capacity and is consistent with the increased protein aggregation known to occur in this cell type. Inhibitory neurons also showed prominent patterns of repression of unfolded protein response pathway (*HSPD1, HSP90AA1, HSPH1*) but additionally displayed signatures of dysregulation of receptor mediated endocytosis pathways (*HSP90AA1, PLCG2, HSPH1*). We further validated the differential expression of neuronal genes using RNAscope *in situ* hybridization on an additional cohort of PD and control brain prefrontal cortex sections for a few downregulated genes, such as *HSP90AA1* and *PLCG2* (Supplementary Fig. 3D) and quantified the RNAscope signals between male PD and control using QuPath (Supplementary Fig. 3E-F).

There was no discernable downregulation pathway signature for microglia, other than phospholipid binding *(LPAR1, ANXA4, AXL)* and virus receptor activity *(CXADR, SELPLG, AXL)*, suggesting microglia play a more active role in initiation of immune responses than negative regulation. Most astrocyte downregulated GO terms consisted of coagulation (*VAV3, LYN, UBASH3B*) indicating its detection of cellular injury. The pattern of T cell downregulated genes was less intuitive, as GO terms were GATOR2 complex (*SESN3, SZT2*), receptor antagonist activity (*MTRNR2L12, MTRNR2L8*) and enhancer binding (*BCH2, MEF2C, LEF1*). Overall, the GO pathway analysis indicates increased gliosis and neuroinflammation in PD with a concomitant repression of neuronal quality control.

### Transcriptional dynamics of excitatory neurons and glial cell subtypes in PD

To infer transcriptional gene expression dynamics of the major identified brain cell types in PD, we applied unsupervised single cell RNA velocity analysis of nascent (unspliced) and mature (spliced) mRNAs to our snRNA-seq dataset using scVelo(*13*), since the sequencing of single nuclei can potentially identify and enrich the amount of unspliced reads from precursor mRNA (pre-mRNA)(*5, 13*). As shown in Supplementary Fig. 2A-B, the single cell RNA velocity fields projected on the Uniform Manifold Approximation and Projection (UMAP)(*14*) plots described the local average velocity and trajectory dynamics of each cell type in PD and healthy control brains. The trajectories of all subpopulations of excitatory neurons were altered, with excitatory neuron subclusters 2 and 3 shifting dramatically, from a left-to-right pattern to a top-to-bottom pattern (Supplementary Fig. 2A-B), indicating a selective transcriptomic signature change for excitatory neuron subpopulations. The genes and pathways that were profoundly altered in excitatory neuron subpopulations were related to neurotransmission (*SYN2, MCTP1, PCLO*), synaptic signaling (*HTR2A, SYN2, CLSTN2, SLC4A10*) and structure (*LRFN5, SLIT1, RAPGEF4*) (Supplementary Fig. 2C-G). Interestingly, a unique pattern of T cell trajectory was found in the PD brains, with enriched *FYN* and *THEMIS* genes, which play a regulatory role in T cell function (Supplementary Fig. 2A-B). In contrast, the dynamics of inhibitory neurons did not show a major pattern change in PD brains.

### Chaperone gene expression patterns in PD correlate with Lewy body pathology in excitatory neurons

We performed phospho-S129-α-synuclein immunohistochemistry on brain prefrontal cortex tissue obtained from the same PD and healthy control individuals used for the snRNA-seq analysis and confirmed the presence of Lewy bodies and Lewy neurites in PD cases with a lack of pathology in controls (Fig. 2A). Quantification of Lewy bodies and Lewy neurites counts in each brain sample showed that PD patients had a range of pathology, but overall, exhibited a significantly higher Lewy body pathology score compared to healthy controls (p = 0.03, unpaired t-test) (Fig. 2B).

**Fig. 2.**
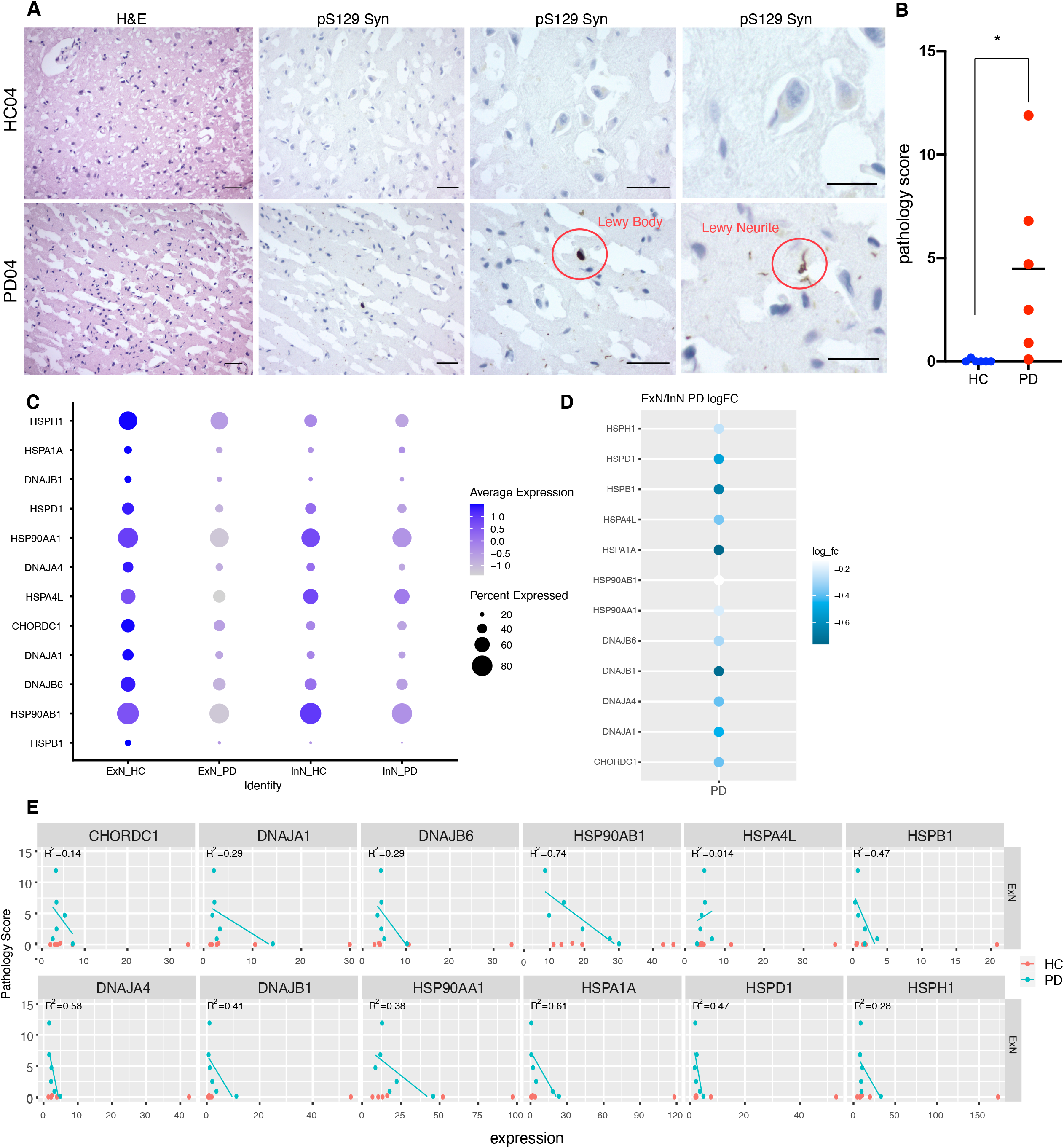
Correlation analysis of Lewy body pathology to chaperone gene expression in PD brains. (**A**) Immunohistochemistry of pS129 α-synuclein (EP1536Y) in formalin fixed prefrontal cortex cortical sections detected α-synuclein-positive Lewy bodies and Lewy neurites. All scale bars represent 50 μm, except in the right-most panel scale bar = 25 μm. (**B**) Bar graph of Lewy body plus Lewy neurite score quantification across conditions (n = 12, unpaired t-test p = 0.03). (**C**) Gene expression dot plot of unfolded protein misfolding pathway in excitatory neurons (ExN)/inhibitory neurons (InN) of PD versus healthy controls (HC). (**D**) Dot plot of gene expression patterns (log_fc: log fold changes) of ExN/InN in PD patients (**E**) Pathology correlation analysis with each protein misfolding gene. Spearman’s correlation coefficient was calculated based on normalized gene expression versus Lewy pathology score.

We sought to correlate the Lewy pathology score with gene expression in excitatory neuron subpopulations, as pathology is largely restricted to these neurons in PD(*15*). We carried out an unbiased correlation analysis and identified postsynaptic GO pathways as the most positively correlated (q = 0.0009), suggesting that the observed synaptic alterations (Fig. 1G) perhaps contribute to synaptic spread of pathology. Protein folding pathways (q = 0.021) were most negatively correlated with the Lewy pathology score, suggesting the decrement of chaperones (Fig. 1H) are likely to increase α-synuclein aggregation and pathology.

Based on our observation that protein folding pathways were significantly downregulated in neurons in PD (Fig. 1H) and that the same pathways were negatively correlated with Lewy pathology scores, we investigated this relationship further. We first plotted the expression levels of significantly downregulated genes in the protein folding pathway in both excitatory and inhibitory neurons. Notably, PD brains showed a lower average expression in most heat shock protein genes in excitatory neurons in comparison to healthy controls (Fig. 2C). While inhibitory neurons also displayed similar trends of downregulation of chaperone genes in PD, it was not as pronounced as the repressive patterns of protein misfolded response genes in excitatory neurons (Fig. 2C). The relative sparing of chaperone expression in inhibitory neurons in PD may contribute to the observation that Lewy bodies are most often found in excitatory neurons but not inhibitory neurons(*15*). Apart from three genes (*HSPH1, HSPB1*, and *CHORDC1*), all misfolded protein response genes exhibit downregulation in excitatory neurons in PD (Fig. 2D). Based on the immunohistochemistry Lewy pathology score (Fig. 2D), we performed a Pearson’s correlation analysis for each gene of the protein folding pathway. Overall, we observed negative correlation patterns, suggesting that pathology is inversely correlated with chaperone gene expression including *HSP90AB1* (R^2^ = 0.74; m = -0.42), *HSPA1A* (R^2^ = 0.61; m = -0.34), and *DNAJA4* (R^2^ = 0.58; m = -2.7) (Fig. 2E). As several of these chaperones have been shown to be directly involved in α-synuclein protein folding, this finding suggests a decrement in chaperone capacity contributes to Lewy body pathology. Interestingly while *HSPA4L* displayed the most prominent difference in gene expression across excitatory and inhibitory neuronal cell types, Lewy pathology showed a minimal correlation (R^2^ = 0.014; m = 0.41), indicating disaggregase (*HSPA4L* encoded Hsp110(*16*), the rate limiting component of the disaggregase) activity are not probably enhanced in PD, adding to the pathology.

### Regulation of PARK and GWAS candidate genes in human PD brains

We corroborated our PD snRNA-seq data by analyzing differential expression of all known 23 PARK genes, including *SNCA* (*PARK1/4*), *PRKN* (*PARK2*), *PINK1* (*PARK6*), *DJ-1* (*PARK7*), and *LRRK2* (*PARK8*), as well as the PD risk gene *GBA* (Fig. 3A-C). *SNCA* and *LRRK2* were upregulated in the nuclei from PD brains, consistent with previous literature that increased expression or activity (in the case of *LRRK2*) is associated with greater disease risk(*17*). Interestingly, *LRRK2* was highly expressed in OPCs and microglia, consistent with previous reports of *LRRK2* expression in monocytes in the blood samples of PD patients(*18*), pointing to its potential role in neuroinflammation. The expression of *PRKN, PINK1, GBA* and *DJ-1* were downregulated in PD brains mainly in excitatory neurons, consistent with a loss-of-function in PD (Fig. 3B). This directionality of expression extends to most PD genes, implying that the modest changes in *PARK* gene expression could contribute to disease risk, as was unequivocally established for *SNCA(19)*. We next sought to correlate PARK gene expression in excitatory neurons with Lewy pathology scores. Interestingly, we found that *LRRK2* (R^2^ = 0.58; m = 1.9), *GIGY2* (*PARK11*) (R^2^ = 0.48; m = 1.6) and *DNAJC13* (*PARK21*) (R^2^ = 0.39; m = 2.7) expression were positively correlated with Lewy pathology scores (Supplementary Fig. 3).

**Fig. 3.**
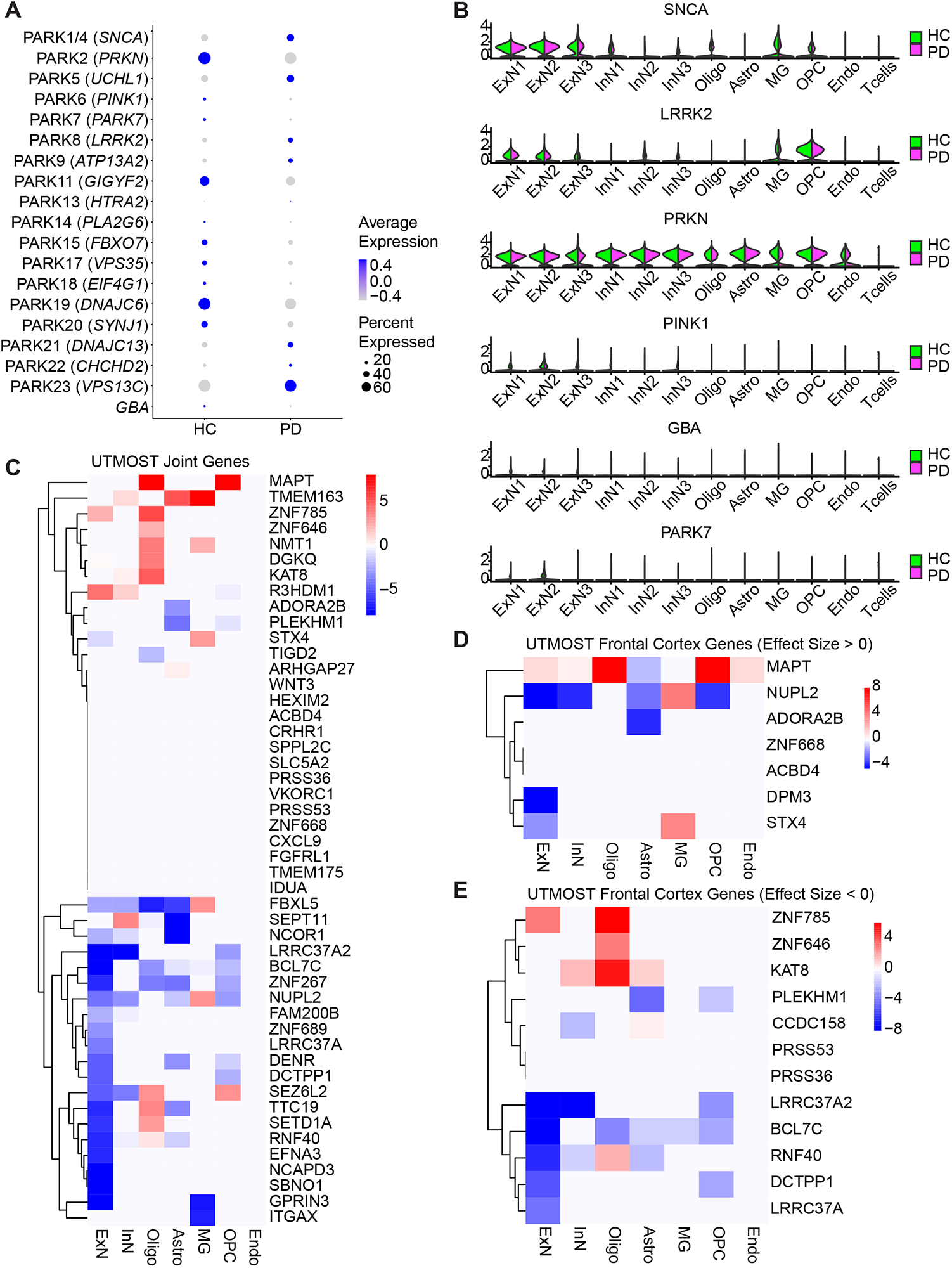
Cell-type-specific regulation of PARK genes and GWAS candidate genes in human PD brains. **(A**) Differential expression of known PARK genes and *GBA* in human PD brains compared to healthy controls (HC). (**B**) Cell-type-specific differential expression of *SNCA, LRRK2, PRNK, PINK1, GBA* and *PARK7* in major neuronal and glial brain cell types in human brains PD versus HC. (**C**) Heatmap for cell-type-specific expression of UTMOST identified most significant genes in PD brains compared to healthy controls. (**D-E**) Heatmap for cell-type-specific differential expression of UTMOST prefrontal cortex most significant genes with effect size greater (D) or less (E) than 0 in human PD brains compared to healthy controls.

Meta-analysis of genome-wide association studies (GWAS) of PD have identified over 50 risk haplotypes for the disease(*20, 21*). To evaluate their expression patterns in human PD brains, we carried out Unified Test for Molecular Signatures (UTMOST) analysis(*22*), using a summary-statistic-based testing framework to quantify the overall gene–trait association. Leveraging data from the NINDS-Genome-Wide Genotyping in Parkinson’s Disease project, we identified 48 genes by the UTMOST multi-tissue joint analysis (Fig. 3C), seven genes in prefrontal cortex (BA9) with a positive effect size and 12 with a negative effect size (Fig. 3D). For each cell type, we plotted the z scores calculated from the t-test of each gene in PD versus healthy controls using the snRNA-seq data. Genes rendered from the joint-tissue analysis can be broadly clustered into a PD upregulated set (Fig. 3D top) and a PD downregulated set (Fig. 3D bottom). Upregulated genes including *MAPT, TMEM163*, and *KAT8* were enriched in oligodendrocytes, while downregulated genes including *LRC37A2, BCL7C*, and *LRRC37A2* were found principally in excitatory neurons. In terms of BA9 specific genes, we observed many overlapping profiles with the joint-tissue gene list. For example, *STX4* and *NUPL2* had positive effect sizes and were highly expressed in microglia (Fig. 3D) whereas *LRRC37A2, BCL7C, RNF40, DCTPP1*, and *LRRC37A*, having negative effect sizes and were downregulated in excitatory neurons (Figs. 1d, 3f). UTMOST analysis underscored the cellular heterogeneity of disease risk genes in PD.

### Cell-cell communications between neurons, glia, and T cells in PD brains

To systematically study the interactions between neurons and other cell types in the brain, we leveraged our human brain snRNA-seq dataset to identify candidate cell-cell communication pathways using CellPhoneDB(*23*), a repository of ligand-receptor interacting pairs. We first curated cell-cell interactions in healthy control brains and then extended the survey of cell-cell communications between cell types in PD brains to enable comparisons. A matrix of the difference of cell-cell interaction numbers in PD brains to healthy controls revealed that overall cell-cell interactions are decreased in PD across all cell types, indicating a general loss of cell-cell communication (Fig. 4A). One of the most dramatic changes in the cell-cell communication is between excitatory neurons and astrocytes, where ∼25% of the interactions were lost in PD compared to healthy controls (Supplementary Table 3). In controls, astrocytes express the EGF receptor which interacts with TGF-β and betacellulin (*BTC*)(*24*) expressed by excitatory neurons. However, these interactions are missing in the PD brains (Fig. 4B). Interestingly, *SIRPA/CD47* signaling is also lost in the PD brains, suggesting an abnormal neuroinflammation process(*25*). Interactions involving activin receptor (*ACVR*) between excitatory neurons and astrocytes were absent along with the interaction between tyrosine kinase *TYRO3* receptor and its ligand *GAS6*. Taken together, the lack of key cell-cell communications between astrocytes and excitatory neurons in the PD brains shows a pattern of abatement of neuron-glia interactions in PD, consistent with an abnormal neuroinflammation process.

**Fig. 4.**
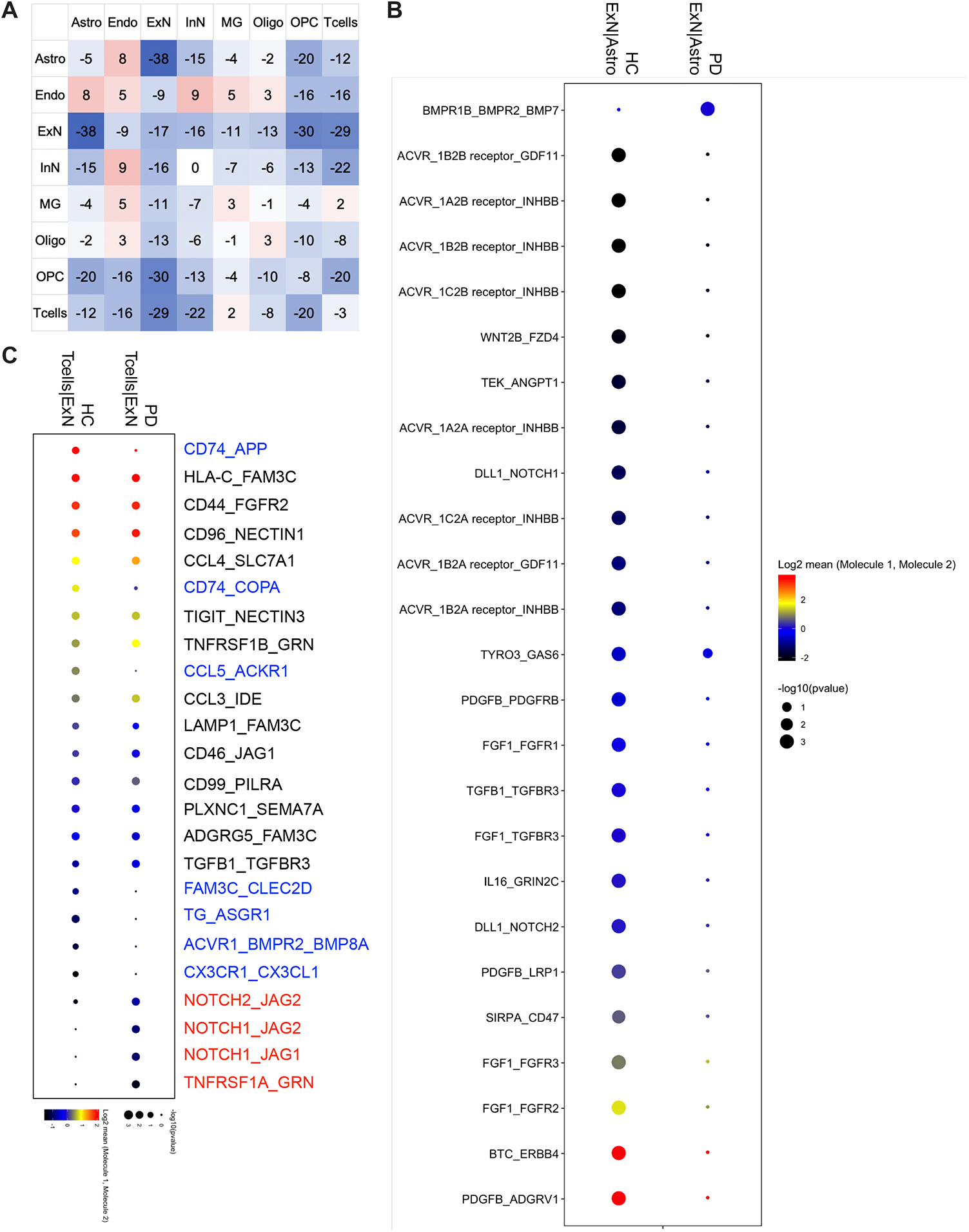
Cell-cell communications between brain cell types in human PD brains. (**A**) Differential matrix of PD (n = 6 individuals) versus healthy controls (HC, n = 6 individuals) cell-cell interaction pair numbers calculated by CellPhoneDB. (**B**) Dot plot of the most differential receptor-ligand interaction pairs between excitatory neurons (ExN) and astrocytes (Astro) in PD and HC brains. (**C**) Dot plot of cell-cell interactions between brain-resident T cells (Tcells) and excitatory neurons (ExN) in PD and HC brains (red: unique interactions in PD; blue: unique interactions in HC).

We previously identified soluble and surface receptor-surface ligand pairs between T-cells and neuronal and glial cells in healthy control brains(*26*). In PD brains, most of the key interactions between T cells and excitatory or inhibitory neurons were retained, including interactions involving the co-inhibitory ligand T cell immunoreceptor with Ig and ITIM domains (*TIGIT*), the brain tissue resident marker of *CD96*, and the chemokines *CCL3* and *CCL4* (Fig. 4C). Additional interaction pairs between the *NOTCH* signaling pathway were observed in PD, indicating a possible pathogenic link between Notch pathway and PD(*27*) (Supplementary Table 3).

### Paired proteomic and transcriptomics analysis of PD brains

Utilizing Label Free Quantitative (LFQ) mass spectrometry, we profiled proteomic changes from the same brain tissues that were used for our snRNA-seq analysis. In total, 35,964 peptides were mapped to 2,534 unique gene symbols. Twenty-two proteins were identified as differentially expressed between PD and healthy controls (two-sided t-test, P < 0.05), with 13 upregulated proteins, including CDH2 (N-Cadherin), GSTT1 (Glutathione-S-transferase theta 1) and N4BP1 (NEDD4 binding protein), as well as nine downregulated proteins, including SEC13, BRCC3, and NDRG2 (Fig. 5A). Most of those proteins show distinct expression patterns across all PD and healthy control brains (Fig. 5B). We analyzed the differentially expressed proteins by GO and show the top upregulated and downregulated GO pathways (Fig. 5C). Top upregulated GO terms include glutathione metabolic process involving glutathione S-transferase (GST), an anti-oxidant enzyme known to protect neurons from reactive oxidative species(*28*). Another upregulated pathway was maintenance of cytoskeleton polarity, which could be indicative of a cellular adaptation to reinforce intracellular protein transport. Downregulated GO pathways included branched chain amino acid (BCAA) metabolic / catabolic processes which is known to affect tyrosine metabolism and dopamine levels(*29*). Also, pathways that regulate COPII vesicle coating and protein transport from ER to Golgi were downregulated, suggesting perturbation in protein transport processes.

**Fig. 5.**
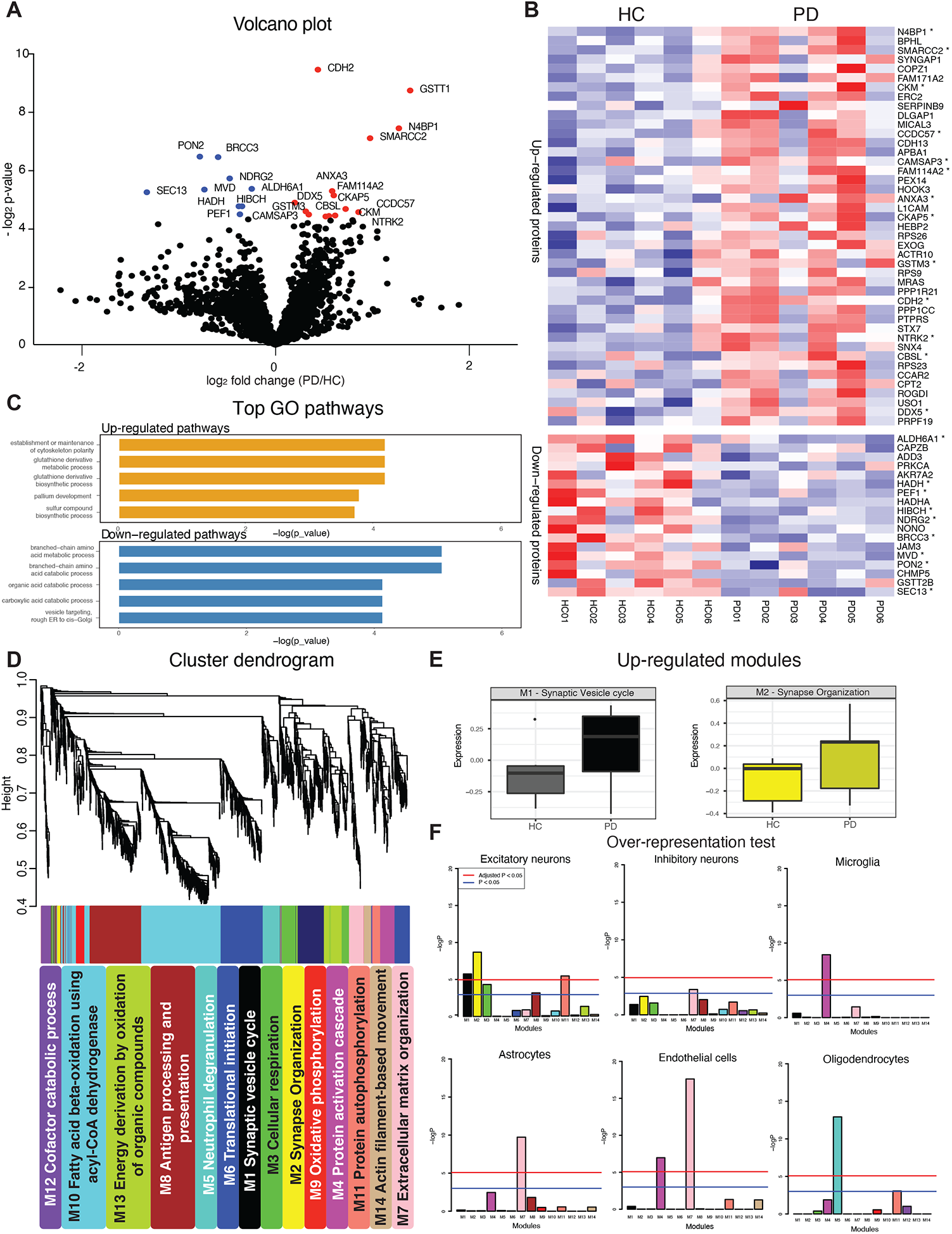
Analyses of the paired proteomics and single cell transcriptomics of PD brains. (**A**) Volcano plot showing significant differentially expressed proteins (n = 2,523 proteins, two-sided t test, P < 0.05) of PD (n = 6 individuals) versus health controls (HC, n = 6 individuals). (**B**) Heat map showing the upregulated (top) and downregulated (bottom) differentially expressed proteins (P < 0.1, asterisked proteins; P < 0.05, row scaled). (**C**) Upregulated (top) and downregulated (bottom) gene ontology (GO) terms (n = 13 upregulated and 9 downregulated genes as input (P < 0.05), hypergeometric test, FDR correction). (**D**) Distinct protein modules (M1-M14) clustered by WGCNA across PD and HC. Each module’s label is the most significantly enriched GO term of that module proteins. (**E**) Eigen-proteins that are highly expressed in PD brains compared to HC in the most upregulated modules of M1 and M2. (**F**) Cell-type enrichment evaluated by module proteins against lists of RNA markers for different brain cell types from the snRNA-seq data using the one-tailed Fisher’s exact test. Blue line: P < 0.05. Red line: Adjusted P < 0.05, Bonferroni correction.

We compared differentially expressed proteins in our proteomic data to their RNA levels in excitatory neurons. As published previously, protein and RNA levels were poorly correlated(*30*). Therefore, we characterized the co-expression pattern in our proteomics data, and performed weighted gene co-expression network analysis (WGCNA) across PD and healthy control samples. We identified 14 modules (M1 to M14). Two of the modules were synapse related (M1 Synaptic vesicle cycle, and M2 Synapse organization), while the remaining modules were associated with GO terms including cellular respiration (M3), protein activation cascade (M4), and antigen processing and presentation (M8) (Fig. 5D). Expression levels of several eigen-proteins were altered in PD compared with the expression levels in healthy control samples. Of note, the two synapse related modules (M1 and M2) were both upregulated in PD (Fig. 5E) consistent with the upregulated pathways in the two neuronal subclusters we found from analysis of the snRNA-seq data (Fig. 1G). To determine whether these modules contain cell-type specific information, we assessed the overlap of the proteins from each module with the cell type marker genes identified from our snRNA-seq data. Accordingly, excitatory neuron-specific RNA markers were enriched in synapse related module M1 (e.g., HPCAL1, NPTX1, and SYN2), M2 (e.g., ERC2, CDH13, and CCK), and M11 (e.g., HPCA, and DCLK1). Interestingly, the enrichment of *SYN2* was also identified using single cell RNA velocity analysis in PD brains (Fig. 3), suggesting a robust enrichment of synaptic signaling at both RNA and protein levels in PD. The M4 module was enriched for microglia markers (PADI2, CD14, and CD163) and endothelial cell markers (ANXA1, and CFH). The M7 module was associated with both astrocyte markers (SLC14A1, AQP4, and AGT) and endothelial cell markers (COL1A2, CAVIN2, and FN1). Oligodendrocyte markers were over-represented in the M5 module, including ANLN, ERMN, and CNP (Fig. 5F). In summary, distinctive differentially expressed proteins and pathways were identified from our proteomics data, and through WCGNA could be attributed to cell types from the snRNAseq data obtained from the same PD patients. A pattern that emerged from this analysis is that synaptic pathways in neurons are impacted in PD.

### Cross-neurodegenerative disease comparisons at both transcriptomic and proteomic levels

To identify shared transcriptomic PD and AD signatures, we compared our snRNA-seq dataset with a landmark AD snRNA-seq dataset which profiled postmortem human brain tissue from the prefrontal cortex (Brodmann area 10, BA10) of 24 AD patients and 24 controls(*8*). We also compared our proteomics data to an analogous proteomics AD dataset of prefrontal cortex(*31*). In the snRNAseq comparisons, we identified 138 DEGs across six cell types, including excitatory neurons, inhibitory neurons, oligodendrocytes, astrocytes, microglia, and oligodendrocytes precursor cells. Overlapping DEGs showed distinct patterns across cell types. Strikingly, there was no PD/AD transcript overlap across neuronal populations. However, there was significant overlap in glial cells in both directions, including oligodendrocytes (upregulated: *SLC38A2, CA2*; downregulated: *UTY, RASGRAF2*), astrocytes (upregulated: *MT2A, MT1E, MRAS, COL27A1, PLEC, M1M, MT1G*; downregulated: *ALDH1A1, BAALC*), and microglia (upregulated: *SPP1, APOE, APOC1, CACNA1A, IRF8*; downregulated: *SYTL3, IFI44L*) (Fig. 6A). We were unable to compare T cell DEGs, as this population was not identified in previous analyses of AD brains(*8*). Mirroring the transcriptomic analyses, the overlapping DE pathways (P < 0.05, FDR) were exclusively found in astrocytes and microglia, with no shared pathways in neuronal cell types (Fig. 6B). The top overlapping pathways for astrocytes were associated with biological processes that involve cadmium or copper ion detoxification, while those for microglia were related to immune responses, cell activation, and protein secretion (Fig. 6C). Collectively, these data suggest commonalities between PD and AD in terms of expression profiling are in non-neuronal cells, particularly glial cells.

**Fig. 6.**
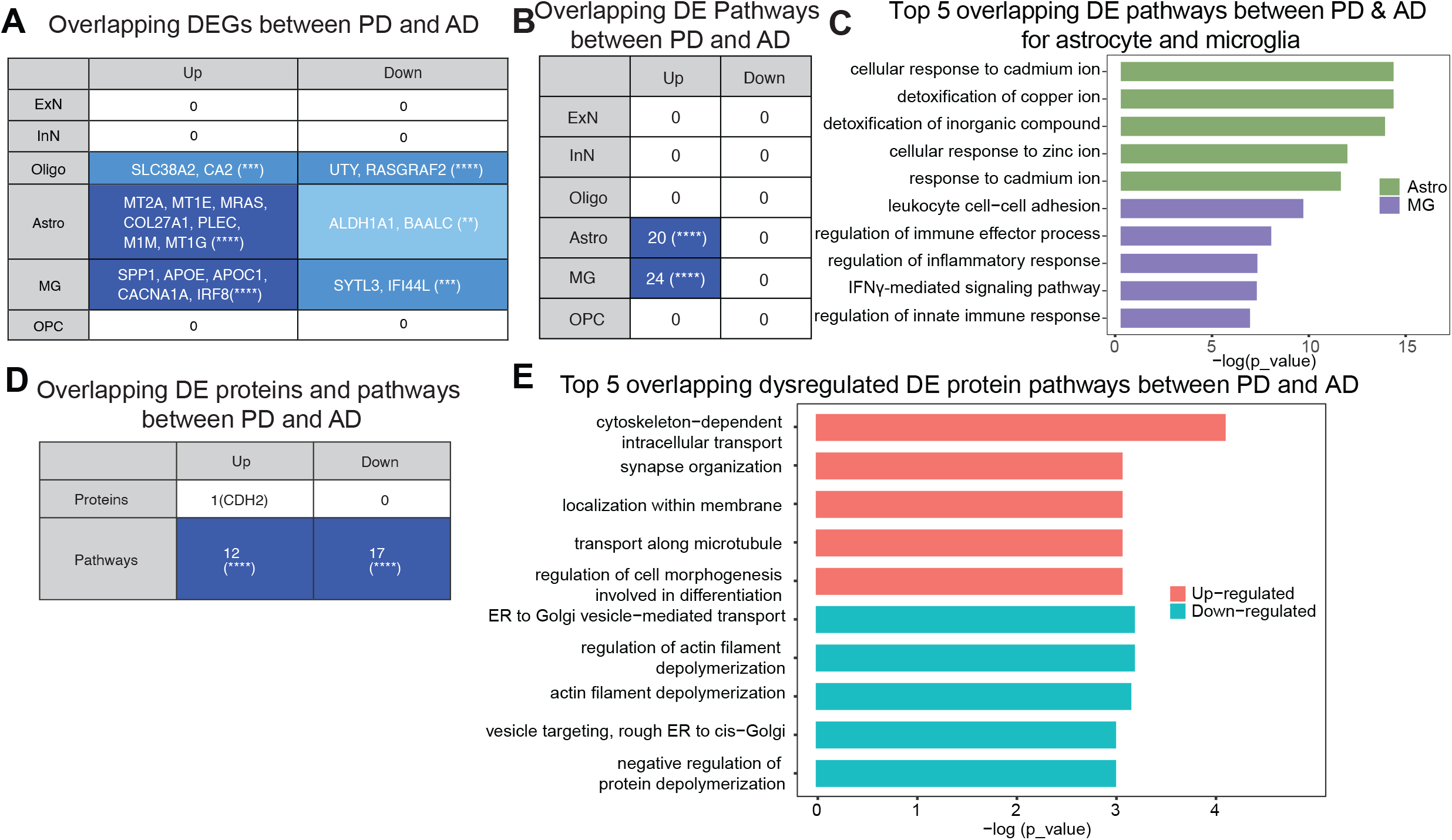
Comparison between PD and AD single cell transcriptomics and proteomics. (**A**) The number of overlapping (over-representation test, (**): P ⩽ 0.01, (***): P ⩽ 0.001, (****): P ⩽ 0.0001) differentially expressed genes (DEGs) for each cell type between PD brain snRNA-seq data and Mathys et al’s AD brain snRNA-seq data (n = 24 AD and 24 healthy controls). (**B**) The number of overlapping (over-representation test, (****): P ⩽ 0.0001) differentially expressed Gene ontology terms (hypergeometric test, FDR correction) for each cell type between PD brain snRNA-seq data and Mathys et al’s AD brain snRNA-seq data. (**C**) Top five overlapping differentially expressed (DE) pathways for astrocytes (Astro) and microglia (MG). (**D**) The number of overlapping (over-representation test, (****): P ⩽ 0.0001) DE proteins (P < 0.05) and pathways (genes with P < 0.1) between PD proteomics and Ping et al’s AD proteomics data (n = 632 upregulated and 1222 downregulated genes for Ping et al’s data). (**E**) Top five overlapping upregulated and downregulated differentially expressed (DE) protein pathways.

We also compared the proteomic profiles in PD and AD in prefrontal cortex. For AD, we used the dataset as reported by Ping et al(*31*) (10 AD and 10 healthy controls; BA9), which was subject to the same mass spectroscopy procedures as for our PD data. We identified 5,143 proteins that were expressed in all samples, with minimal overlap between the diseases. We found overlap for 12 upregulated pathways and 17 downregulated pathways (Fig. 6D). The top upregulated pathways are linked to cytoskeleton dependent intracellular transport, whereas downregulated pathways are associated with ER to Golgi vesicle transport as well as filament depolymerization (Fig. 6E), akin to what we found in Fig. 5C. Thus, both at the protein and RNA levels, AD and PD appear to initiate distinct disease processes.

## DISCUSSION

While several underlying mechanisms have been suggested for the pathogenesis of PD, there have not been effective treatments that prevent disease progression. Thus, establishing the unbiased molecular pathology of the disease is critical to generate hypotheses that can lead to therapeutic interventions that will more definitively elucidate the major pathogenic insult leading to PD. With rapid advancement of single cell RNA sequencing technologies and computational methods, it is possible to identify detailed disease signatures in PD at a single-cell resolution, while the currently published studies(*32*) of postmortem single-cell transcriptomic studies are limited in scope. Here we provide an extensive analysis profiling nearly 80,000 brain nuclei from prefrontal cortex of late-stage PD brains. We demonstrate that α-synuclein pathology is inversely correlated with chaperone expression in excitatory neurons. We further extended our analysis by utilizing advanced computational methods including UTMOST, scVelo, CellPhoneDB, and WGCNA to gain a comprehensive understanding of the dataset. By examining cell-cell interactions, we found a selective abatement of neuron-astrocyte interactions with enhanced neuroinflammation. Finally, in comparing AD to PD pre-frontal cortex, we did not find common expressed genes in neurons but identified many shared differentially expressed genes in glial cells, suggesting that disease etiology in PD and AD are distinct. We augmented the study with proteomic analysis and cross-comparisons with AD datasets of the same brain region providing valuable insights into the pathways of neurodegeneration. Thus, we provide a deep definition of the underlying molecular pathology for PD.

### Cell-type specific pathways impacted in PD

Using snRNA-seq, we profiled unique disease signatures of neuronal and glial cell populations correlating them with GWAS and proteomic data. GO pathway analysis of the single-cell transcriptomics data (Fig. 1G-H) identified cell-type specific disrupted biological processes, with distinct pathways highlighted in the major cell types of the brain: neurons (upregulated: synapse assembly, synaptic organization; downregulated: protein folding), astrocytes (upregulated: response to metal ion, downregulated: coagulation), microglia (upregulated: immunological activation: downregulated: phospholipid binding), oligodendrocytes/OPC (upregulated: regulation of cell growth) and T cells (upregulated: T cell differentiation).

We observed upregulation of presynaptic assembly/organization pathways by DEG analysis (Fig. 1G) and RNA velocity in neuronal populations. Several genes (*EPHA6, NLGN4Y, CBLN2*) encode for synaptic cell-adhesion molecules that are known to orchestrate synapse formation and maintenance. Overexpression of these pathways may reflect perturbations in synaptic architecture that is crucial to maintaining basal circuitry in the prefrontal cortex. These results are congruent with the idea that cognitive impairments in PD are mainly attributed to altered activity of the neural pathways connecting the anterior striatum with the prefrontal cortical region(*33*). Single cell RNA velocity results (Supplementary Fig. 2) underscored these synaptic changes in excitatory neurons. WGCNA modules (M1: synaptic vesicle cycling, M2: synapse organization) derived from proteomic interrogation of prefrontal cortex further support the concept that synaptic pathways are defective in PD excitatory neurons. Downregulated pathways in PD neurons were all related to protein folding and will be discussed further below.

The number of oligodendrocytes was found to be decreased in PD patient brains, while the number of microglia was increased compared to healthy controls, suggesting that glial cell populations play differential, but a significant role in PD pathology (Fig. 1). Similar trends have been described in the analysis of a PD midbrain dataset(*34*). Also, significantly increased proportion of microglia has been found in a recent single cell study of anterior cingulate cortex across Lewy body diseases(*35*). Many of the DEGs in oligodendrocytes and OPCs, encode GWAS hits such as *MAPT* and *TMEM163* (Fig. 3). Microglia are the most abundant immune cells in the CNS and participate in PD disease progression(*36*). In our dataset, microglia had the greatest number of DEGs (135 upregulated, 178 downregulated) with a distinct signature of gliosis in the up-regulated GO pathways. Genes responsible for myeloid cell differentiation (*FOS, STAT3, RUNX1*) and regulation of hemopoiesis (*BCL6, IL4R, JAK3*) were activated and are strong indicators of neuroinflammation in the prefrontal cortex. In the case of astrocytes, even though we did not observe any changes in numbers, we observed clear upregulation of pathways in astrocytes related to stress responses to heavy metals. Astrocytes are known to protect neurons by accumulating heavy metals commensurate with their deposition in PD(*32*).

One notable finding was the identification of T cells in PD brains. In this population, there was gene enrichment for pathways regulating homeostasis of cell numbers (*IL7R, IL15, MALT1*) and regulation of T cell differentiation (*CAMK4, CD28, ZAP70*). Our results are congruent with recent findings that clonal expansion of α-synuclein reactive T cells have been identified in the brains of PD patients(*11, 12*) and that CD4^+^ T cells contribute to neurodegeneration in Lewy body dementia(*37*). Our findings add to existing data suggesting that T cell mediated immunity plays an important role in PD and α-synucleinopathies. Our results also indicate the need to carefully profile T cells in in PD at different stages, other movement disorders as well as AD and tauopathies for comparative and mechanistic purposes.

### Cellular correlates of Lewy body pathology

We observed severe repression of protein folding and misfolded protein response pathways (*HSP90AB1, HSPB1, DNAJB6*) in both excitatory and inhibitory neuronal populations, though the effect was more pronounced in excitatory neurons (Fig. 2C, D). Due to the constant proteostatic stress in synaptic terminals, impairment of chaperone machinery has been known to cause functional deficits in neurons, which has long-term impact on cortical neurotransmission(*38*). We evaluated the effects of proteostasis deficits by correlating chaperone/heat shock protein gene expression in excitatory neurons with phospho-S129-α-synuclein positive Lewy pathology, revealing negative associations of chaperone expression in relation to Lewy bodies and Lewy neurites (Fig. 2E). These data underscore the utility of leveraging single cell/nucleus data derived from postmortem human brain tissue in the context of neuropathology.

Most of the heat shock proteins that are encoded by genes in the downregulated protein folding pathways are neuroprotective since they directly interfere with α-synuclein misfolding. For example, Hsp90ab1, which is highly negatively correlated with Lewy pathology score, is known to inhibit α-synuclein aggregation by interacting with oligomeric species(*39*). Another example is heat shock protein DNAJB6 which has been shown to suppress α-synuclein aggregation as knockout models resulted in higher number of exogenous preformed fibrils (PFFs)-induced synuclein aggregates(*40*). DNAJB1 in conjunction with the HSPA1A (Hsp70) disaggregase-complex is linked to disassembling α-synuclein amyloid fibrils *in vitro*(*41*). Moreover, it has been shown that expression of heat shock proteins such as DNAJB1 in gut microbiome can cross-react with heat shock proteins expressed in stressed tissue in autoimmune disorders, suggesting a novel mechanism of gut-brain interactions potentially mediating PD by T cells activated in the gut(*42*). Finally, our observations support that the molecular chaperones whose expression is decreased in PD serve active roles in α-synuclein protein quality control have the potential to become novel therapeutic targets for PD.

### Common pathways in AD and PD

In a cross-comparison of our PD dataset with the AD dataset generated by Mathys et al.(*8*), we did not observe overlapping genes or pathways in excitatory or inhibitory neuronal populations. In addition, we did not identify overlapping genes and pathways in oligodendrocytes and OPCs, even though these cell types appear to have a prominent role in AD(*8*). Rather, we observed upregulation of common pathways within astrocyte and microglia populations. The top upregulated pathways in astrocytes in PD are stress responses to heavy metals. These detoxification pathways are also observed in AD, suggesting that a common disease signature in astrocytes.

The top five overlapping pathways between PD and AD for microglia consisted of pathways for interferon gamma signaling and innate immune cell regulation, implying a shared immune response across the two neurodegenerative diseases. One of the overlapping genes was *ApoE* which encodes for apolipoprotein E and is a well-studied genetic risk factor for AD, dementia with Lewy Bodies, and Parkinson disease dementia(*43, 44*). *ApoE* was downregulated in both PD and AD in microglia. Studies have shown activated microglia promotes tau pathogenesis in an *ApoE* dependent manner(*45*). Our transcriptomic profiling of PD patients highlights a significant role of ApoE in microglia and calls for further investigation on how ApoE promotes neuroinflammation. It also underscores while PD and AD share microglial neuroinflammatory signatures, T cell mediated adaptive immunity may be different between AD, PD and other α-synucleinopathies such as Lewy body dementia(*37*).

In sum, the tandem employment of single nucleus transcriptomic studies and unbiased proteomic interrogation of the prefrontal cortex in PD provides invaluable insight into the complex molecular and cellular pathobiology of late-stage PD. Taken together, these datasets are a valuable resource to the scientific research community for assessment of the underlying pathogenesis of PD as well as a tool to evaluate overarching neurodegenerative disorder dysfunctions through cross comparisons with analogous datasets obtained from diseases including movement disorders with cognitive decline (e.g., corticobasal degeneration and progressive supranuclear palsy), the AD spectrum, and tauopathies for a better mechanistic understand of disease processes as well as provide viable targets for therapeutic intervention.

## MATERIALS AND METHODS

### Postmortem human brain tissue cohort

Prefrontal cortex (Brodmann area 9, Dorsolateral prefrontal cortex) obtained from postmortem PD and non-diseased age- and sex-matched control individuals was provided by brain tissue biorepository at Nathan Kline Institute and New York University (NKI/NYUGSOM)(*46, 47*). Study participants were allocated into disease or control groups based on overall Parkinson’s disease clinical diagnosis and brain pathology as described previously(*2*)^,(*48*),^(*49*). Dissected brain tissues were fresh frozen samples stored at -80 °C until use.

### Nuclei isolation from postmortem frozen human brain tissue

Nuclei were isolated from postmortem human brain tissue as previously described(*26*) with modifications. All procedures were carried out on ice or at 4 °C. In brief, fresh frozen tissue (50 to 100 mg) was homogenized in 15 ml of ice-cold nuclei homogenization buffer [2 M sucrose, 10 mM Hepes (pH 7.9), 25 mM KCl, 1 mM EDTA (pH 8.0), 10% glycerol, and ribonuclease (RNase) inhibitors freshly added (80 U/ml)] using a 15-ml Wheaton Dounce tissue grinder (10 strokes with the loose pestle and 10 strokes with the tight pestle). The homogenate was transferred into a 50 ml ultracentrifuge tube on top of 10 ml of fresh nuclei homogenization buffer cushion and centrifuged at 24,000 rpm for 60 min at 4 °C on ultracentrifuge. The supernatant was removed, and the pellet was resuspended in 1 ml of nuclei resuspension buffer [15 mM Hepes (pH 7.5), 15 mM NaCl, 60 mM KCl, 2 mM MgCl_2_, 3 mM CaCl_2_, and RNase inhibitors freshly added (80 U/ml)] and counted on a hemocytometer with Trypan Blue staining. The nuclei were centrifuged at 800g for 10 min at 4°C with a swing bucket adaptor and resuspended at a concentration of 500 to 1000 nuclei/ul in the nuclei resuspension buffer for the next step of 10x Genomics Chromium loading.

### Droplet-based single nucleus RNA sequencing

The snRNA-seq libraries were prepared by the Chromium Single Cell 3′ Reagent Kit v3 chemistry according to the manufacturer’s instructions (10x Genomics). The generated snRNA-seq libraries were sequenced using Illumina NovaSeq6000 S4 at a sequencing depth of 300 million reads per sample.

### snRNA-seq data alignment

For snRNA-seq of brain tissues, a custom pre-mRNA human genome reference was generated with human genome reference GRCh38_3.0.0 (available from 10x Genomics) that included pre-mRNA sequences, and snRNA-seq data were aligned to this GRCh38–pre-mRNA reference to map both unspliced pre-mRNA and mature mRNA using CellRanger version 3.1.0.

### Single cell quality control, clustering and cell type annotation

After quality control filtering of eliminating nuclei with less than 200 genes (poor quality nuclei) and more than 7,000 genes (potential doublets) per nucleus, we profiled 77,384 brain nuclei and detected a median of 2,598 genes and 5,639 transcripts per nucleus (Supplementary Fig. 1D, 3027 mean genes and 8629 mean transcripts per nucleus). Seurat (version 4.0.2) single cell analysis R package was used for processing the snRNA-seq data. Basically, the top 2000 most variable genes across all nuclei in each sample were identified, followed by the integration of all samples and dimensionality reduction using principal components analysis (PCA). Then, Uniform Manifold Approximation and Projection for Dimension Reduction (UMAP) was applied to visualize all cell clusters, and the classification and annotation of distinct cell types was based on known marker genes of each major brain cell type and the entire single nucleus gene expression matrix (Supplementary Fig. 1G).

### Differential expression (DE) analysis

Differential expression analysis for snRNA-seq data was performed using the Wilcoxon Rank Sum test using the function FindMarkers of the Seurat package (3.2.3) in R. For T cells, the threshold was set as the absolute value of the expression log fold change of PD over healthy controls being larger than 0.75 with no other constraints. For other cell types, the genes were filtered using the default parameters. We followed the same procedure as that by Mathys et al(*8*), and Grubman et al(*50*). in AD snRNA-seq data analysis, and the parameters were set to the default values for all the cell types. As for differential analysis for proteomics data, we used the two-sided t test for both our data and Ping et al(*51*) AD data. Proteins were called differentially expressed if P < 0.05.

### Gene ontology (GO) pathway analysis

Gene-set and protein enrichment analysis was performed using the function enrichGO from the R package clusterProfiler. GO terms from biological process (BP), cellular component (CC), and molecular function (MF) subontologies were considered. The background genes were set to be all the protein coding genes in the snRNA-seq data. We used the default values for the other parameters. For plotting, we prioritized BP over CC, followed by MF.

### Single cell RNA velocity (scVelo) analysis of RNA velocity and gene identification

The RNA velocity was computed using the Python package scVelo. The PD samples and the HC samples were processed separately to better compare the differences between these two conditions. First, the function filter_and_normalize was applied to conduct data filtering, normalization and log transformation. Next, for each cell, the first- and second-order moments were calculated by the function pp.moments. Then, the gene velocities and graph were calculated, and the velocities were projected to the precomputed UMAP plot using velocity, velocity_graph, and velocity_embedding_stream functions respectively, with default parameters. And finally, to identify cell type specific genes that are responsible for the inferred velocities compared to all the other cell types, the function rank_velocity_genes was employed for each cell type separately.

### Phospho-α-synuclein staining

Immunocytochemistry and histopathology were performed on the prefrontal cortex obtained from the same cohort of brains as snRNA-seq and proteomics. Slabs of frozen prefrontal cortex that remained after sequencing was fixed in 10% neutral buffered formalin overnight at room temperature and then processed into paraffin blocks at the Yale Pathology Tissue Services Histology Core. Hematoxylin and Eosin (H&E) staining and immunohistochemical staining for phospho-serine-129 (pS129) α-Synuclein (Abcam; ab51253) were performed according to manufacturer’s protocol / product datasheet, and with appropriate controls. Quantitative scoring of synuclein pathology was performed in a blinded manner. Lewy neurite (LN) score was defined as the average number of pS129 positive Lewy neurites counted in 10 consecutive high power fields (400x magnification). Similarly, the Lewy body score was defined as the average number of pS129 positive Lewy bodies counted in 10 consecutive high power fields. The combined quantitative Lewy pathology score was defined as the sum of the LN score and Lewy body score weighted by a factor of 10 (LN+10* Lewy body) in the prefrontal cortex. LN scores in the prefrontal cortex were also subdivided into semiquantitative scoring categories, as follows: “None” was defined LN score of zero; “Mild (+)” was defined as LN score > 0 to 2; “Moderate (++)” was defined as LN score > 2 to 5; and “Severe (+++)” was defined as LN score > 5.

### RNAscope *in situ* hybridization, imaging, and quantification

To validate gene expression differences identified by snRNA-seq, RNA *in situ* hybridization was performed using the RNAscope Multiplex Fluorescent v2 Assay (Advanced Cell Diagnostics, Inc., Newark, CA, USA). Human brain tissue from the prefrontal cortex of either PD or controls was flash frozen and stored at -80°C. Prior to performing the assay, frozen tissue was embedded in OCT and sectioned at 10μm using cryostat. RNA *in situ* hybridization was performed according to the manufacturer’s protocol. Briefly, fresh frozen sections were fixed in 10% NBF for 1 hour at room temperature, then sequentially dehydrated in 50% EtOH, 70% EtOH, 100% EtOH, and 100% EtOH for 5 minutes each at room temperature. Tissue sections were pretreated with hydrogen peroxide and protease to block endogenous peroxidase activity and optimally permeabilize the sections. Target RNAscope probes, such as *EPHA6, CBLN2, PLCG2* and *HSP90AA1*, were hybridized to tissue sections, signal was amplified, and fluorescent dyes of Opal Dye 570 or 650 were applied. Positive control RNAscope probes for human POLR2S, PPIB, UBC, and HPRT-1 and a negative control probe for *Bacillus subtilis* DapB were used to ensure detection of target RNA signals and a lack of non-specific probe binding. All tissue sections were counterstained with DAPI. Images were captured using a Leica SP8 motorized staging confocal microscope with a 20x lens (Leica #506517), while confocal imaging settings were kept constant between healthy controls and PD. RNA expression was quantified using QuPath, by detecting the cells using DAPI channel, identifying and measuring probe channel signals, and calculating estimated spot count per cell for the entire cryosection. RNA expression in human brain sections from PD patients was compared to that in healthy control human brains in approximately 10,000 nuclei from each tissue section, and statistical significance was determined using Student’s t-test.

### Single cell-level annotation of post-GWAS candidate genes

To determine if any PD related GWAS genes were also DEGs in a particular cell type of our snRNA-seq data, we downloaded the single nucleotide polymorphism (SNP) summary statistics from the NINDS-Genome-Wide Genotyping in Parkinson’s Disease Project (https://www.ncbi.nlm.nih.gov/projects/gap/cgi-bin/study.cgi?study_id=phs000089.v3.p2) to impute the cross-tissue expression and the overall gene-trait association using the Python package UTMOST. The resulting genes that are associated with PD in prefrontal cortex (BA9) and the overall joint-tissues were extracted and filtered to select genes having P value < 0.05 after Bonferroni correction. Next, conditional analysis was performed using the function conditional_test_geneset to mitigate gene co-regulation or linkage disequilibrium (LD). Finally, the z scores were calculated from the two-sided t test of PD versus healthy controls for each cell type in the snRNA-seq data using the two sets of the UTMOST PD-associated genes.

### Label Free Quantitative mass spectrometry analysis

LFQ sample preparation was carried out similar to that described in Henderson et al., 2015(*52*), but with slight updated modification. Brain tissue of control and PD samples were first dissolved in 400μL RIPA buffer containing protease with Phosphatase inhibitor cocktail and sonicated at 10% amplitude for 15 seconds with one second burst. Sonicated samples were then centrifuged at 4 °C for 10 minutes at 14.6K RPM on a bench top centrifuge. 100μL of supernatant was transferred to a new tube for protein extraction with Chloroform:Methanol:Water method. Protein pellet was air dried, dissolved in 80-μl 8 M urea containing 0.4 M ammonium bicarbonate. Protein amount was determined by NanoDrop and 50μg total protein amount was aliquoted and brought up to 80ul with 8 M urea containing 0.4 M ammonium bicarbonate. Samples were then reduced with DTT for 10 minutes at 37 °C, and Cysteine were alkylated with iodoacetamide at room temperature for 30 minutes in the dark. 10 μL of 0.1 μg/μL LysC was then added for enzymatic digestion overnight at 37 °C; followed by that addition of 2 μL of 0.5 μg/μL trypsin with incubation at 37 °C for 6 hours. The digestion was quenched with 0.1% trifluoroacetic acid (TFA) then desalted utilizing a C18 UltraMicro Spin column (The Nest Group). The effluents from the de-salting step were dried and re-dissolved in 5 μL 70 % FA and 35 μL 0.1 % TFA. An aliquot was taken to obtain total digested protein amount. A 1:10 dilution of Pierce Retention Time CalibrationMixture (Cat# 88321) was added to each sample prior to injecting onto the UPLC coupled Orbitrap Fusion mass spectrometer system for normalization of the LFQ data.

Data Dependent Acquisition (DDA) LC MS/MS data collection for LFQ was performed on a Thermo Scientific Orbitrap Fusion connected to a Waters nanoACQUITY UPLC system equipped with a Waters Symmetry® C18 180 μm × 20 mm trap column and a 1.7-μm, 75 μm × 250 mm nanoACQUITY UPLC column (35 °C). The digests were diluted to 0.05 μg/μL with 0.1 % TFA prior to injecting 5 μL of each triplicate analysis in block randomized order. To ensure a high level of identification and quantitation integrity, a resolution of 120,000 and 30,000 was utilized for MS and MS/MS data collection, respectively. MS and MS/MS (from Higher-energy C-Trap Dissociation (HCD)) spectra were acquired using a three second cycle time with Dynamic Exclusion on. All MS (Profile) and MS/MS (centroid) peaks were detected in the Orbitrap. Trapping was carried out for 3 min at 5 μl/min in 99 % Buffer A (0.1 % FA in water) and 1 % Buffer B [(0.075 % FA in acetonitrile (ACN)] prior to eluting with linear gradients that reach 25 % B at 150 min, 50 % B at 170 min, and 85 % B at 175 min; then back down to 3% at 182 min.. Two blanks (1st 100 % ACN, 2nd Buffer A) followed each injection to ensure there was no sample carry over. The LC–MS/MS LFQ data was processed with Progenesis QI software (Nonlinear Dynamics, version.4.2) with protein identification carried out using the Mascot search algorithm (Matrix Science, v. 2.7). The Progenesis QI software performs feature/peptide extraction, chromatographic/spectral alignment (one run was chosen as a reference for alignment), data filtering, and quantitation of peptides and proteins. A normalization factor for each run was calculated to account for differences in sample load between injections as well as differences in ionization. The normalization factor was determined by comparing the abundance of the spike in Pierce Retention Time Calibration mixture among all the samples. The experimental design was set up to group multiple injections from each run. The algorithm then tabulated raw and normalized abundances, and maximum fold change for each feature in the data set. The combined MS/MS spectra were exported as .mgf (Mascot generic files) for database searching. The Mascot search results were exported as .xml files using a significance cutoff of p < 0.05 and FDR of 1 % and then imported into the Progenesis QI software, where search hits were assigned to corresponding aligned spectral features. Relative protein-level fold changes were calculated from the sum of all unique and non-conflicting, normalized peptide ion abundances for each protein on each run. Additional downstream biostatistical analyses were conducted utilizing a custom R script.

### Weighted gene co-expression network analysis (WGCNA) module detection on the proteomics dataset

The function blockwiseModules in the R package WGCNA was used to identify modules of highly correlated proteins across PD and healthy control samples. Module eigen-protein was calculated as the first principal component of each of the resulting 13 protein modules and was used to summarize and represent the expression pattern of all the proteins in that module. The label for each module was chosen to be the top significant Gene Ontology (GO) terms using the function enrichGO.

### Over-representation analyses for protein module cell-type RNA marker enrichment

Over-representation test was performed using the one-tailed Fisher’s exact test with the R function phyper to compare each WGCNA protein module against the RNA markers for each cell type from the snRNA-seq data. Bonferroni adjusted P values were calculated to adjust for multiple comparisons. The background for the over-representation test was set as the number of genes in each RNA marker list.

## Supporting information

Supplementary Table 1 Postmortem Brain Cases

Supplementary Table 2 Differentially Expressed Genes

Supplementary Table 3 CellPhoneDB Analysis

## Acknowledgments

This work was supported by an Aligning Science Across Parkinson’s (ASAP; D.A.H., S.S.C., L.Z.), NIH RF1 NS110354 (S.S.C.), Yale/NIDA Neuroproteomic Center (DA018343-11A1). We thank Jesse Cedarbaum and Philip Coish for reading manuscript. The authors also acknowledge Yale Pathology Tissue Services for assistance in tissue processing and staining and Yale Center for Genome Analysis for 10x Genomics library preparation and Illumina sequencing. We also thank Florine Collins and Jean Kanyo for proteomics sample preparation and data collection. The necessary mass spectrometers and the accompany biotechnology tools within the MS & Proteomics Resource at Yale University is funded in part by the Yale School of Medicine and by the Office of The Director, National Institutes of Health (S10OD02365101A1, S10OD019967, and S10OD018034), while additional support came from NIH Yale/NIDA Neuroproteomics Centre (DA018343). The study is funded by the joint efforts of the Michael J. Fox Foundation for Parkinson’s Research (MJFF) and the Aligning Science Across Parkinson’s (ASAP) initiative. MJFF administers the grant [ASAP-000529] on behalf of ASAP and itself. For the purpose of open access, the author has applied a CC-BY public copyright license to the Author Accepted Manuscript (AAM) version arising from this submission.

## Author contributions

D.A.H., S.S.C., and L.Z. conceptualized the study. L.Z. performed single nucleus RNA sequencing experiments, genome alignment, and single cell quality control and clustering analysis. T.L.L. performed mass spectrometry experiments. P.P.G. performed pathology assay and analysis. S.C. and I.H. performed RNAscope, imaging and quantification. S.D.G. provided all human brain specimens. B.Z. and L.Z. performed computational analysis with assistance from J.P., J.W., C.S. and H.Z. B.Z., J.P., D.A.H., S.S.C. and L.Z. wrote the manuscript with input from all authors.

## Competing interests

Authors declare that they have no competing interests.

## Data and materials availability

All postmortem brain single nucleus RNA sequencing data will be available through the National Center of Biotechnology Information’s Gene Expression Omnibus (GEO) at an accession number, including raw sequencing data of fastq files and processed data of gene expression matrix. Raw mass spectrometry data have been deposited in the PRIDE repository and will be made publicly available. Additional data related to this paper may be requested from the authors. Code for Seurat, UTMOST, scVelo, CellPhoneDB, and WGCNA is available at github.com. Additional code is available upon request.

## Notes

### Competing Interest Statement

The authors have declared no competing interest.

