## Supplementary Material for "Single-cell transcriptomic and proteomic analysis of Parkinson’s disease Brains"

**Supplementary Materials**

Fig. S1. Single nucleus transcriptomics quality control of human brain prefrontal cortex in PD.

Fig. S2. Single cell RNA velocity field describes transcriptional dynamics of neuronal and glial cell types in PD brain.

Fig. S3. RNAscope in situ hybridization validation of differentially expressed genes in PD and healthy control brains.

Table S1. Postmortem Brain Cases.

Table S2. Differentially Expressed Genes.

Table S3. CellPhoneDB Analysis.

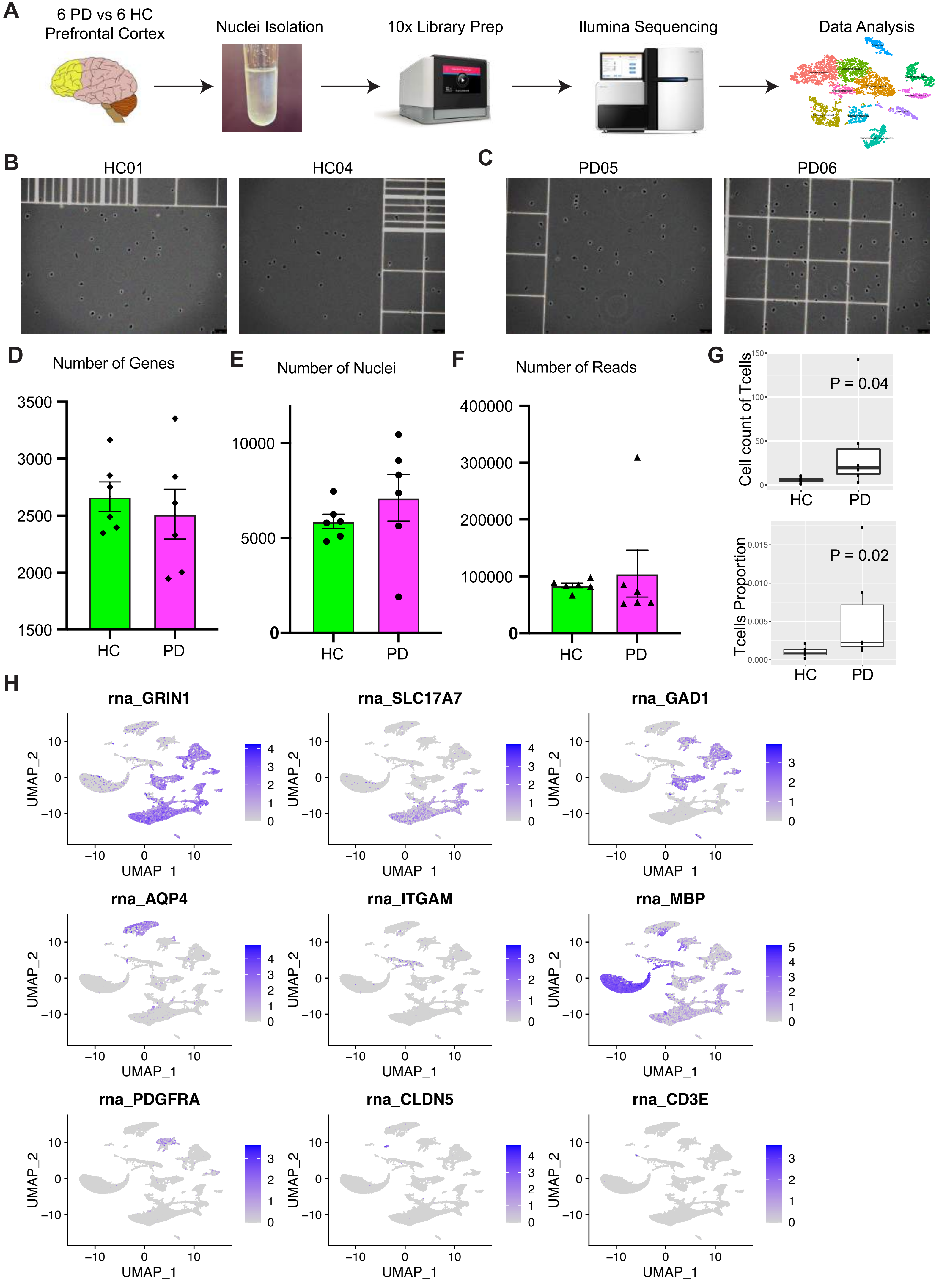
A
6 PD vs 6 HC Prefrontal Cortex
Nuclei Isolation
10x Library Prep
Ilumina Sequencing
Data Analysis
B
HC01
HC04
C
PD05
PD06
D
Number of Genes
E
Number of Nuclei
F
Number of Reads
G
Cell count of Tcells
P = 0.04
Tcells Proportion
P = 0.02
HC
PD
H
rna_GRIN1
rna_SLC17A7
rna_GAD1
rna_AQP4
rna_ITGAM
rna_MBP
rna_PDGFRA
rna_CLDN5
rna_CD3E
UMAP_1
UMAP_2

Figure S1

**Fig. S1. Single nucleus transcriptomics quality control of human brain prefrontal cortex in PD.** (**A**) Single nucleus RNA sequencing workflow of Parkinson's disease (PD) human postmortem brain prefrontal cortex. (**B-C**) Bright filed images of single nucleus suspension (sucrose gradient ultracentrifugation isolated human brain nuclei) in healthy controls (HC) (B) and PD (C). (**D-F**) Single nucleus RNA sequencing quality controls of number of genes per nucleus in each individual (D), number of nuclei per individual (E), and number of reads per nucleus in each individual (F) (n = 6 PD and 6 HC). (**G**) Box plots of cell counts of brain-resident T cells (top panel, P = 0.04) and T cells proportion (bottom panel, P = 0.02) in PD or healthy controls (HC) with Wilcoxon Rank-Sum test. (**H**) UMAP plotting of brain cell type marker gene expression in the postmortem human brains of PD and HC (n = 77,384 nuclei, 6 PD and 6 HC). *GRIN1*, neurons; *SLC17A7*, excitatory neurons; *GAD1*, inhibitory neurons; *AQP4*, astrocytes; *ITGAM*, microglia; *MBP*, oligodendrocytes; *PDGFRA*, oligodendrocyte precursor cells; *CLDN5*, endothelial cells; *CD3E*, brain-resident T cells.

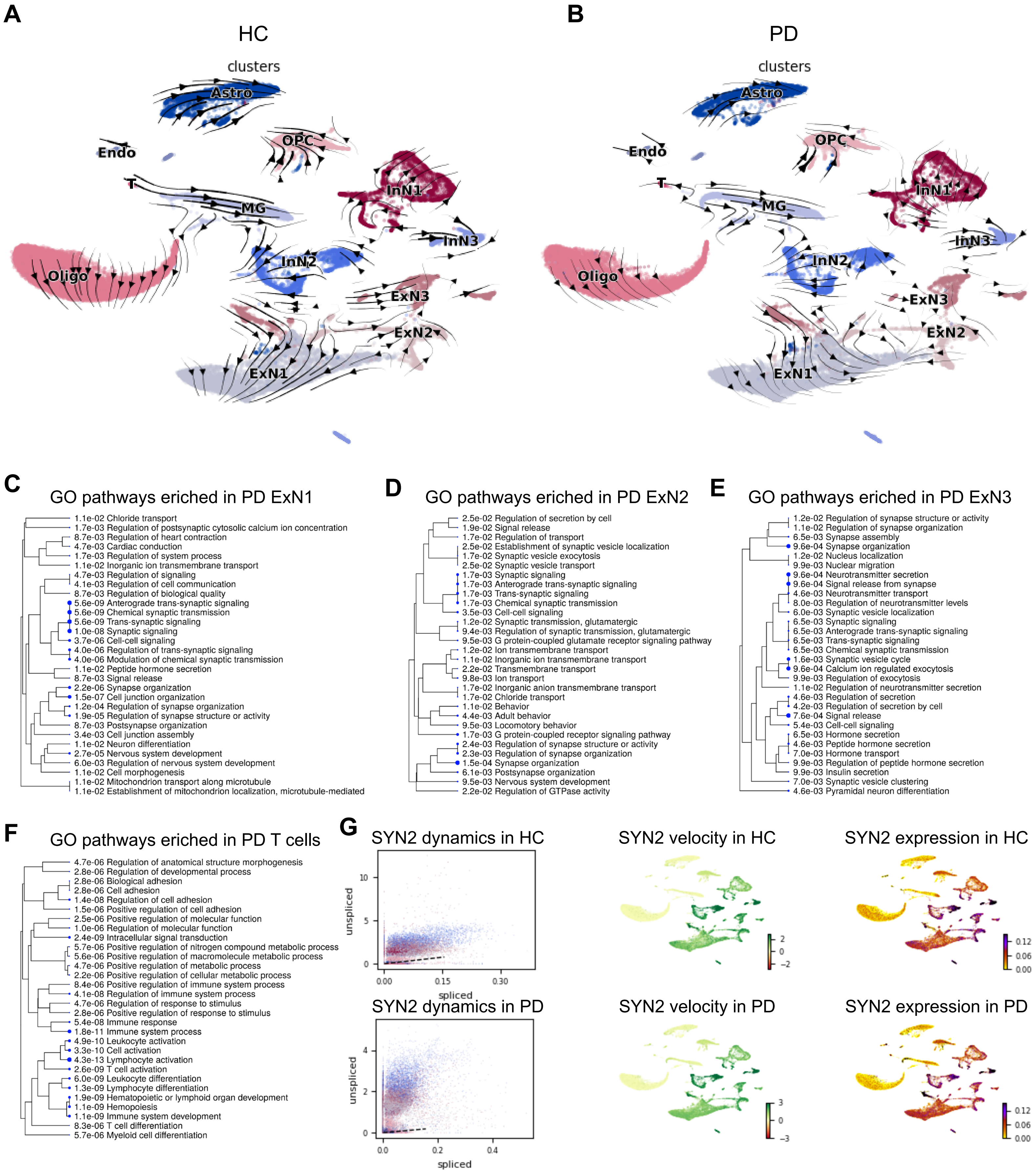
A
HC
clusters
Astro
Endo
OPC
T
MG
InN1
InN3
Oligo
InN2
ExN3
ExN2
ExN1
B
PD
clusters
Astro
Endo
OPC
T
MG
InN1
InN3
Oligo
InN2
ExN3
ExN2
ExN1
C
GO pathways eriched in PD ExN1
1.1e-02 Chloride transport
1.7e-03 Regulation of postsynaptic cytosolic calcium ion concentration
8.7e-03 Regulation of heart contraction
4.7e-03 Cardiac conduction
1.7e-03 Regulation of system process
1.1e-02 Inorganic ion transmembrane transport
4.7e-03 Regulation of signaling
4.1e-03 Regulation of cell communication
8.7e-03 Regulation of biological quality
5.6e-09 Anterograde trans-synaptic signaling
5.6e-09 Chemical synaptic transmission
5.6e-09 Trans-synaptic signaling
1.0e-08 Synaptic signaling
3.7e-06 Cell-cell signaling
4.0e-06 Regulation of trans-synaptic signaling
4.0e-06 Modulation of chemical synaptic transmission
1.1e-02 Peptide hormone secretion
8.7e-03 Signal release
2.2e-06 Synapse organization
1.5e-07 Cell junction organization
1.2e-04 Regulation of synapse organization
1.9e-05 Regulation of synapse structure or activity
8.7e-03 Postsynapse organization
3.4e-03 Cell junction assembly
1.1e-02 Neuron differentiation
2.7e-05 Nervous system development
6.0e-03 Regulation of nervous system development
1.1e-02 Cell morphogenesis
1.1e-02 Mitochondrion transport along microtubule
1.1e-02 Establishment of mitochondrion localization, microtubule-mediated
D
GO pathways eriched in PD ExN2
2.5e-02 Regulation of secretion by cell
1.9e-02 Signal release
1.7e-02 Regulation of transport
2.5e-02 Establishment of synaptic vesicle localization
1.7e-02 Synaptic vesicle exocytosis
2.5e-02 Synaptic vesicle transport
1.7e-03 Synaptic signaling
1.7e-03 Anterograde trans-synaptic signaling
1.7e-03 Trans-synaptic signaling
1.7e-03 Chemical synaptic transmission
3.5e-03 Cell-cell signaling
1.2e-02 Synaptic transmission, glutamatergic
9.4e-03 Regulation of synaptic transmission, glutamatergic
9.5e-03 G protein-coupled glutamate receptor signaling pathway
1.2e-02 Ion transmembrane transport
1.1e-02 Inorganic ion transmembrane transport
2.2e-02 Transmembrane transport
9.8e-03 Ion transport
1.7e-02 Inorganic anion transmembrane transport
1.7e-02 Chloride transport
1.1e-02 Behavior
4.4e-03 Adult behavior
9.5e-03 Locomotory behavior
1.7e-03 G protein-coupled receptor signaling pathway
2.4e-03 Regulation of synapse structure or activity
2.3e-03 Regulation of synapse organization
1.5e-04 Synapse organization
6.1e-03 Postsynapse organization
9.5e-03 Nervous system development
2.2e-02 Regulation of GTPase activity
E
GO pathways eriched in PD ExN3
1.2e-02 Regulation of synapse structure or activity
1.1e-02 Regulation of synapse organization
6.5e-03 Synapse assembly
9.6e-04 Synapse organization
1.2e-02 Nucleus localization
9.9e-03 Nuclear migration
9.6e-04 Neurotransmitter secretion
9.6e-04 Signal release from synapse
4.6e-03 Neurotransmitter transport
8.0e-03 Regulation of neurotransmitter levels
6.0e-03 Synaptic vesicle localization
6.5e-03 Synaptic signaling
6.5e-03 Anterograde trans-synaptic signaling
6.5e-03 Trans-synaptic signaling
6.5e-03 Chemical synaptic transmission
1.6e-03 Synaptic vesicle cycle
9.6e-04 Calcium ion regulated exocytosis
9.9e-03 Regulation of exocytosis
1.1e-02 Regulation of neurotransmitter secretion
4.6e-03 Regulation of secretion
4.2e-03 Regulation of secretion by cell
7.6e-04 Signal release
5.4e-03 Cell-cell signaling
6.5e-03 Hormone secretion
4.6e-03 Peptide hormone secretion
7.0e-03 Hormone transport
9.9e-03 Regulation of peptide hormone secretion
9.9e-03 Insulin secretion
7.0e-03 Synaptic vesicle clustering
4.6e-03 Pyramidal neuron differentiation
F
GO pathways eriched in PD T cells
4.7e-06 Regulation of anatomical structure morphogenesis
2.8e-06 Regulation of developmental process
2.8e-06 Biological adhesion
2.8e-06 Cell adhesion
1.4e-08 Regulation of cell adhesion
1.5e-06 Positive regulation of cell adhesion
2.5e-06 Positive regulation of molecular function
1.0e-06 Regulation of molecular function
2.4e-09 Intracellular signal transduction
5.7e-06 Positive regulation of nitrogen compound metabolic process
5.6e-06 Positive regulation of macromolecule metabolic process
4.7e-06 Positive regulation of metabolic process
2.2e-06 Positive regulation of cellular metabolic process
8.4e-06 Positive regulation of immune system process
4.1e-08 Regulation of immune system process
4.7e-06 Regulation of response to stimulus
2.8e-06 Positive regulation of response to stimulus
5.4e-08 Immune response
1.8e-11 Immune system process
4.9e-10 Leukocyte activation
3.3e-10 Cell activation
4.3e-13 Lymphocyte activation
2.6e-09 T cell activation
6.0e-09 Leukocyte differentiation
1.3e-09 Lymphocyte differentiation
1.9e-09 Hematopoietic or lymphoid organ development
1.1e-09 Hemopoiesis
1.1e-09 Immune system development
8.3e-06 T cell differentiation
5.7e-06 Myeloid cell differentiation
G
SYN2 dynamics in HC
unspliced
spliced
SYN2 velocity in HC
SYN2 expression in HC
SYN2 dynamics in PD
unspliced
spliced
SYN2 velocity in PD
SYN2 expression in PD

Figure S2

**Fig. S2. Single cell RNA velocity field describes transcriptional dynamics of neuronal and glial cell types in PD brain.** (**A-B**) Single cell RNA velocity filed projected on to UMAP plot of either healthy control (HC) (A) or Parkinson's disease (PD) (B) human brains. Arrows show the local average velocity evaluated on a regular grid. (**C-F**) Gene ontology (GO) pathway analysis of PD enriched genes identified by RNA velocity analysis in excitatory neuron subcluster 1 (ExN1) (C), excitatory neuron subcluster 2 (ExN2) (D), excitatory neuron subcluster 3 (ExN3) (E) and brain-resident T cells (F). (**G**) Transcriptional dynamics, velocity and expression of *SYN2* in HC or PD.

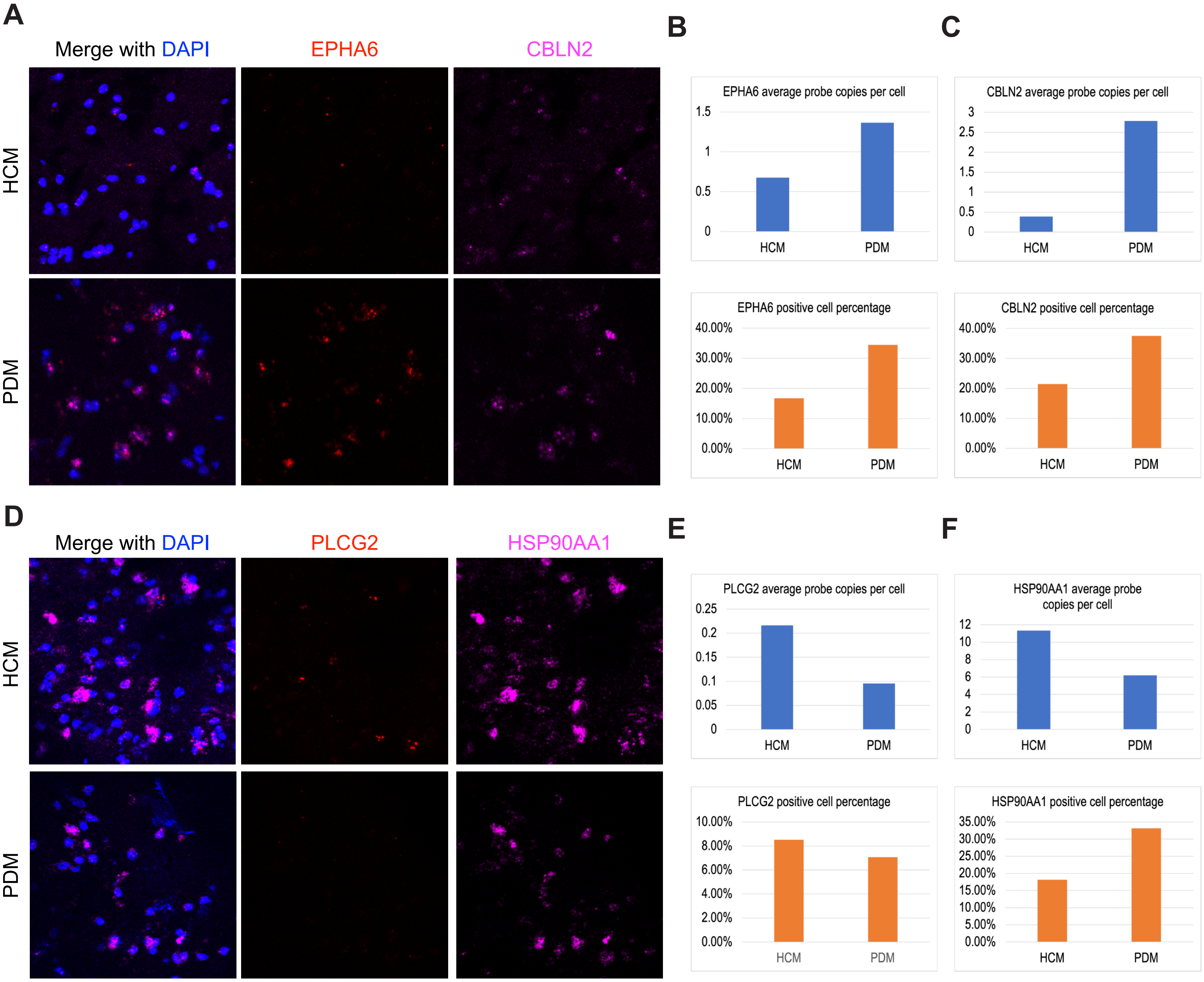
A
Merge with DAPI
EPHA6
CBLN2
HCM
PDM
B
EPHA6 average probe copies per cell
1.5
1
0.5
0
HCM
PDM
EPHA6 positive cell percentage
40.00%
30.00%
20.00%
10.00%
0.00%
HCM
PDM
C
CBLN2 average probe copies per cell
3
2.5
2
1.5
1
0.5
0
HCM
PDM
CBLN2 positive cell percentage
40.00%
30.00%
20.00%
10.00%
0.00%
HCM
PDM
D
Merge with DAPI
PLCG2
HSP90AA1
HCM
PDM
E
PLCG2 average probe copies per cell
0.25
0.2
0.15
0.1
0.05
0
HCM
PDM
PLCG2 positive cell percentage
10.00%
8.00%
6.00%
4.00%
2.00%
0.00%
HCM
PDM
F
HSP90AA1 average probe copies per cell
12
10
8
6
4
2
0
HCM
PDM
HSP90AA1 positive cell percentage
35.00%
30.00%
25.00%
20.00%
15.00%
10.00%
5.00%
0.00%
HCM
PDM

Figure S3

**Fig. S3. RNAscope *in situ* hybridization validation of differentially expressed genes in PD and healthy control brains.** (**A**) RNAscope images of *EPHA6* and *CBLN2* in male Parkinson's disease (PDM) or male healthy control (HCM) brains. Blue: DAPI; Red: *EPHA6*; Magenta: *CBLN2*. (**B-C**) QuPath quantification of RNAscope images for *EPHA6* (B) and *CBLN2* (C) (HCM, n = 13,219 cells; PDM, n = 8,963 cells). Upper panels: average probe copies per cell; Lower panels: positive cell percentage in all cells. (**D**) RNAscope images of *PLCG2* and *HSP90AA1* in male Parkinson's disease (PDM) or male healthy control (HCM) brains. Blue: DAPI; Red: *PLCG2*; Magenta: *HSP90AA1*. (**E-F**) QuPath quantification of RNAscope images for *PLCG2* (E) and *HSP90AA1* (F) (HCM, n = 16,071 cells; PDM, n = 11,543 cells). Upper panels: average probe copies per cell; Lower panels: positive cell percentage in all cells.
